# In silico structure-based analysis of the predicted protein-protein interaction of Syntaxin-18, a putative receptor of *Peregrinus maidis* Ashmead (Hemiptera: *Delphacidae*) with Maize mosaic virus glycoprotein

**DOI:** 10.1101/2022.02.02.478912

**Authors:** Melvin A. Castrosanto, Apel Jae N. Clemente, Anna E. Whitfield, Karen B. Alviar

## Abstract

The corn planthopper, *Peregrinus maidis*, is a widely distributed insect pest which serves as a vector of two phytopathogenic viruses, Maize mosaic virus (MMV) and Maize stripe virus (MStV). It transmits the viruses in a persistent and propagative manner. MMV is an alphanucleorhabdovirus with a negative-sense, single-stranded (ss) RNA unsegmented genome. One identified insect vector protein that may serve as receptor to MMV is Syntaxin-18 (PmStx18) which belongs to the SNAREs (soluble N-ethylmaleimide-sensitive factor attachment protein receptors). SNAREs play major roles in the final stage of docking and subsequent fusion of diverse vesicle-mediated transport events. In this work, in silico analysis of the interaction of MMV glycoprotein (MMV G) and PmStx18 was performed. Various freely available protein-protein docking web servers were used to predict the 3D complex of MMV G and PmStx18. Analysis and protein-protein interaction (PPI) count showed that the complex predicted by the ZDOCK server has the highest number of interaction and highest affinity, as suggested by the calculated solvation free energy gain upon formation of the interface (Δ^i^G = −31 kcal/mol). Molecular dynamics simulation of the complex revealed important interactions at the interface over the course of 50 ns. This is the first in silico analysis performed for the interaction on a putative receptor of *P. maidis* and MMV G. The results of the protein-protein interaction prediction provide novel information for studying the role of STX18 in the transport, docking and fusion events involved in virus particle transport in the insect vector cells and its release.

## INTRODUCTION

*Peregrinus maidis* (Ashmead), commonly known as corn planthopper, is just one of the pests of corn devastating the maize-producing regions mostly in tropical and subtropical areas.. It is a known vector of two disease-causing plant pathogens of corn, the *Maize mosaic virus* (MMV) and the *Maize stripe virus* (MStV) (Jourdan-Ruf et al., 1995). Once *P. maidis* acquires MMV, the virus persistsand replicates in the insect reaching a threshold level in the adult stage (Barandoc-Alviar et al., 2017). MMV is the causal agent of the mosaic disease of corn broadly occurring in Africa, Asia, and in the Americas (Centre for Agriculture and Bioscience International (CABI). The loss of yield due to corn diseases in the United States and Ontario, Canada accounts for $76.51 USD per acre in 2012 to 2015 (Mueller et al., 2016). Additionally, an annual loss of $ 480 million dollars in the sub-Saharan Africa was reported by Karavina (2014) due to the incidence of the streak disease of maize. Similarly, a study by Kannan et al. Kannan et al. (2018) reported that there is 70% loss in the global yield of the commodity since 1920 due to *Maize dwarf mosaic virus* (MDMV). In the Philippines, recent visits in corn fields in CALABARZON and the Bicol region show *P. maidis* devastation due to insect damage and visible symptoms of MMV disease in majority of the crops surveyed.

Identification of *P. maidis* receptors for MMV G through high throughput membrane yeast two hybrid system revealed a putative interacting protein known as Syntaxin-18 ((Alviar et al., 2022)). This protein belongs to the Syntaxin family, classified under soluble N-ethylmaleimide-sensitive factor-attachment protein receptor (SNARE) which is important in membrane fusion (Fasshauer et al., 1998). Specifically, the proteins in this family play roles in vesicle docking and/or fusion within exocytic as well as endocytic pathways and are principally located in the endoplasmic reticulum (Hatsuzawa et al., 2000). Moreover, SNARE proteins could either be vesicle membrane SNAREs (v-SNAREs) or target membrane SNAREs (t-SNAREs) (Yoon & Munson, 2018). Syntaxin-18 (STX18) is a t-SNARE protein (Bossis et al., 2005; UniProtKB).

Protein protein interaction (PPI) is involved in various biological processes such as cell-to-cell interactions, and metabolic and developmental control (Rao et al., 2014). Targeting PPIs has recently become a strategy in drug development due to their association with diseases (Lu et al., 2020). Several research papers on prediction of PPIs between protein sequences usually employ in silico analysis which utilizes methods such as sequence-based and structure-based approaches. In this paper, the 3D structure of MMV G and PmStx18 were modeled and refined. Then, the protein-protein interaction of MMV G and PmStx18 was explored and the interacting residues at the interface were identified. Molecular dynamics simulation was conducted to monitor the stability of the predicted complex.

## MATERIALS AND METHODS

### Homology modelling of PmStx18 and MMV G

The protein sequence of MMV G was retrieved from the protein database of NCBI. Both the protein sequences of PmStx18 and MMV G were submitted to I-TASSER (Iterative Threading Assembly Refinement) server found (Yang et al., 2015) for the prediction of the proteins’ secondary and tertiary structure as well as the binding sites.

### Model validation and refinement

The initial homology models (MMV G and PmStx18) were validated using the SWISS-MODEL structure assessment tool, noting the MolProbity score, Ramachandran favored, Ramachandran outliers, and rotamer outliers (Artimo et al., 2012). Then, refinement was done via GalaxyWEB GalaxyRefine tool (Ko et al., 2012) to improve the quality of the models. GalaxyWEB is a web server for protein structure prediction, refinement, and related methods developed by the Computational Biology Lab, Department of Chemistry, Seoul National University (Ko et al., 2012). The refined models were again validated and compared to the initial models.

### Protein-Protein Docking

The possible interaction between MMV G and PmStx18 were predicted through protein-protein docking using three different webservers – PatchDock (Schneidman-Duhovny et al., 2005), ZDOCK (Pierce et al., 2014), HADDOCK (De Vries et al., 2010), InterEvDock2 (Quignot et al., 2018), pyDockWEB (Jiménez-García et al., 2013), LZerD (Christoffer et al., 2021), Vakser Lab (Tovchigrechko & Vakser, 2006), ClusPro 2.0 (Kozakov et al., 2017), and HDOCK (Yan et al., 2020). The best model in all protein-protein docking web servers were refined using GalaxyWEB GalaxyRefineComplex (Heo et al., 2016). Then, the refined complexes were subjected to PDBePISA (Krissinel & Henrick, 2007) analysis, where the interactions at the interface and the solvation free energy gain upon the formation of the interface (Δ^i^G) were calculated. The MMV G-PmStx18 docking pose having a more negative Δ^i^G and more interaction at the interface was visualized and was chosen to undergo molecular dynamics simulation.

### Molecular dynamics simulation

To obtain the thermally equilibrated system of the MMV G-PmStx18 complex, it was subjected to MD simulation using Desmond (Bowers et al., 2006). The complex was solvated with water molecules using the SPC model in the Desmond System Builder tool. An orthorhombic simulation box shape with distances of 4.0 Å x 4.0 Å x 10.0 Å was generated. The system was neutralized by adding Na+ and Cl- and 0.15 M salt concentration to conserve isosmotic condition. The simulation (NPT) was set to 50 ns with a recording interval of 50 ps at 300 K and 1.01325 bar. Comparison of the initial and final frame of the simulation was done by overlapping the structures. Distance of initial hydrogen bonding interactions were also monitored over the course of the simulation.

## RESULTS AND DISCUSSION

The *Maize mosaic alphanucleorhabdovirus* (MMV) which is vectored by a delphacid planthopper, *Peregrinus maidis*, is the known causative agent of mosaic disease of corn. Advancements in management and control approaches have been developed throughout the years where most of them now are designed to work at the molecular level. A study by Yao et al. (2013) presented an RNAi-based gene knockdown in *P. maidis* by targeting essential genes through oral delivery and microinjection of ATPase B and V-ATPase D double-stranded RNA (dsRNA). Similarly, another molecular protocol has been introduced by Klobasa et al. (2021) which employs CRISPR/CAS9 genome editing in *P. maidis* embryos as basis for gene silencing and germline transformation.

An important prerequisite of development of molecular-based methods in management of corn pests and diseases relies on the knowledge of the interaction of vector and pathogen. It has been previously reported that plant rhabdovirus glycoprotein spikes are predicted to interact with receptors in the midgut allowing the entry of virions into the epithelial cells (Dietzgen et al., 2016). However, there is still limited knowledge on the mechanism of interaction of the receptor protein of *P. maidis* and MMV glycoprotein (MMV G) which may explain the mechanism of infection, replication, and intercellular dissemination of viral particles within the insect host. In connection with this, in silico analysis was carried out in this work to investigate the putative t-SNARE STX18 of *P. maidis* (PmStx18) and its interaction with MMV G. It is assumed that this interaction has a significant role in mediating the infection of MMV within its insect vector host.

### Homology Modelling of PmStx18 and MMV G

Both protein sequences of PmStx18 and MMV G were submitted to I-TASSER to generate the three-dimensional models. Predicted 3D models are provided with C-score, TM-score and RMSD. In I-TASSER, the confidence score (C-score) is calculated based on the significance of threading template alignments with values ranging from [-5, 2] where a C-score greater than −1.5 signifies a model of correct topology (Zhang, 2008). Both the template modeling score (TM-score) and root mean square deviation (RMSD) are known standards for measuring the accuracy of structure modeling. A TM-score greater than 0.5 indicates similar topology between two predicted structures and a TM-score which is less than or equal to 0.17 indicates similarity between two randomly selected structure from the PDB library (Zhang, 2008). The average distance of all residue pairs between two structures is measured by RMSD while TM-score measures structural similarity. According to Roy et al. (2010), predicted structures with 1–2 Å RMSD are high-resolution models which are generated from close homologous template. Structures with RMSD of 2~5 Å are medium-resolution models generated from threading of distantly homologous templates but can still be used for identification of functionally important residues (Roy et al., 2010). Furthermore, the TM-score function is the proposed scale to solve the problem with RMSD since the latter is sensitive to local error while the former is independent of the protein length, thus, template aligned regions may have better quality due to fewer residues than full-length model (Zhang & Skolnick, 2005).

PmStx18 and MMV G structures were not chosen based on the rank provided by I-TASSER but rather based on the C-score values considering the C-score cutoff of −1.5. Figure 1 shows the I-TASSER predicted structures of PmStx18 and MMV G. Based on the C-score cutoff, both the generated models of PmStx18 and MMV G have the best quality among the given models as indicated by their C-score values of −1.11 and −1.32, respectively. Moreover, PmStx18 has an RMSD of 7.4± 4.2Å while MMV G has 15.4± 3.4Å. Although the two structures may not be as accurate due to their high RMSD values, it could still be considered that the predicted structure for PmStx18 is of correct global topology since its calculated TM-score of 0.58± 0.14 is greater than the cutoff value, indicating good structure. For MMV G, its TM-score of 0.37± 0.13 may not be greater than the cutoff value, however, it is significantly close to 0.5 and does not indicate random similarity.

**Figure 1.**
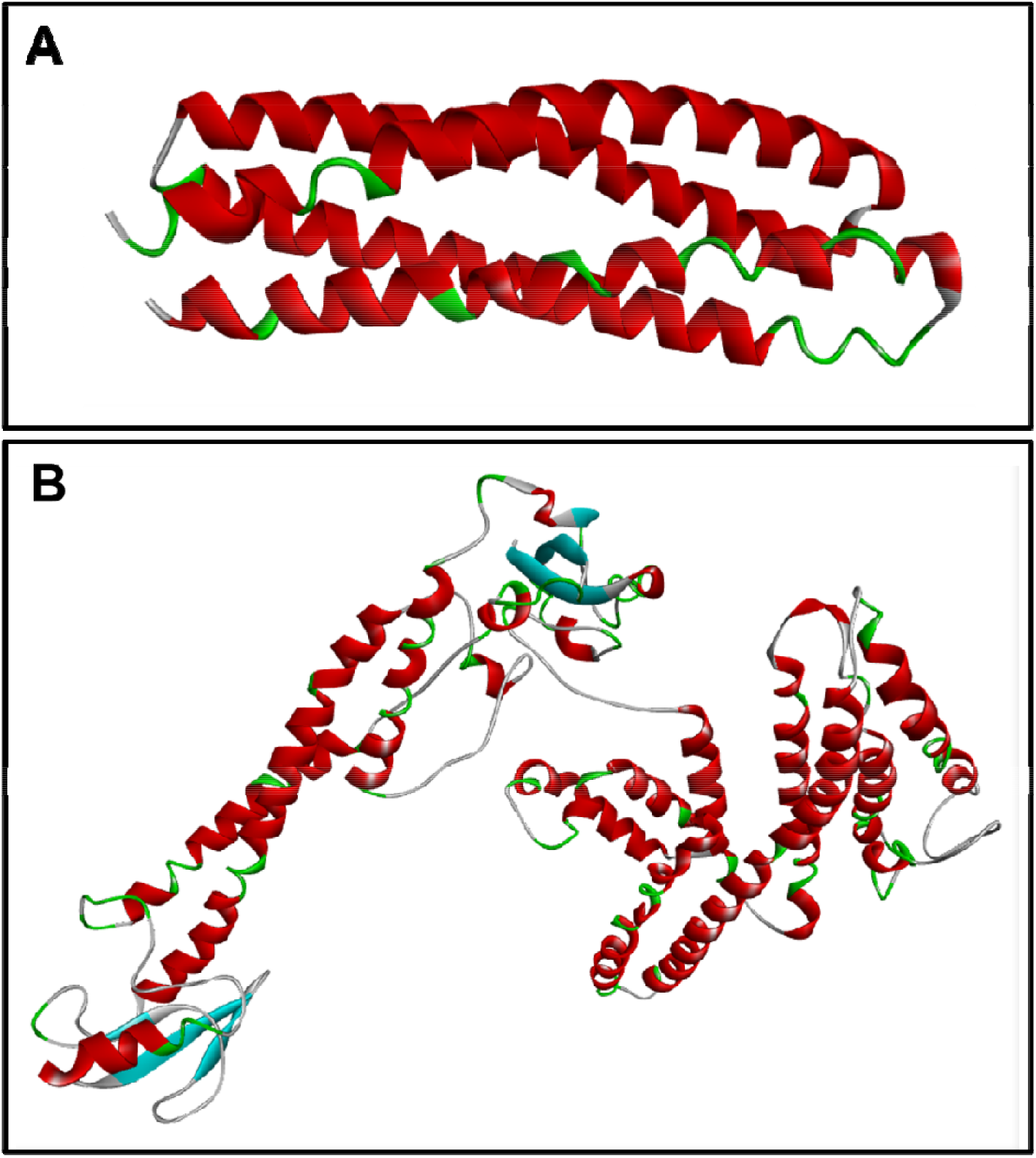
Three-dimensional models of PmStx18 (A) and MMV G (B) predicted in I-TASSER (helix – red; blue – strand; green/gray – turn/coils).

A confidence score ranging from 0 −9 were also provided to indicate the confidence of the predicted secondary structure. From the prediction, PmStx18 have 54 coils and 116 helices whereas the latter is found mostly at positions 61-140. Most of the helices were scored with 8 and 9 while the coils have scores ranging from 0-6 (Supplemental figure—Appendix 1). Majority of MMV G are mostly coils with 307 residues, while helices and strands are only 147 and 137, respectively. Most of the helices are found at positions 300 to 340, as well as in between 520 to 580 having confidence scores which mostly range form 7-9. Additionally, coils are mostly found at positions 20-60, 120-280 and 440-520 while the strands are scattered in the sequence (Supplemental figure—Appendix 2).

### Model validation and refinement

The initial models (MMV G and PmStx18) generated by the I-TASSER web server needs significant structural improvement as suggested by the high percentage of Ramachandran outlier residues (Figure 2 – left plots). Red dots in the plot indicates individual residues. Those residues lying on the white region represents the outliers and needs to be corrected. On the other hand, residues lying on a darker green region implies that their positioning, as well as their stereochemistry are favored. After subjecting to structural refinement using the GalaxyRefine of GalaxyWEB server, which performed repeated structure perturbation and subsequent overall structural relaxation by molecular dynamics simulation, improvement of the Ramachandran plots was evident as more of the residues clumped on the darker green regions. Moreover, the lowering of MolProbity score and rotamer outliers (Table 1) for both MMV G and PmStx18 models signifies a better protein model. MolProbity provides a score that is based on the model quality at both the global and local levels (Chen et al., 2010). A lower score indicates better model quality. The correctness of the sidechain prediction is characterized by the rotamer outliers – having a low outlier means a better model.

**Table 1.**
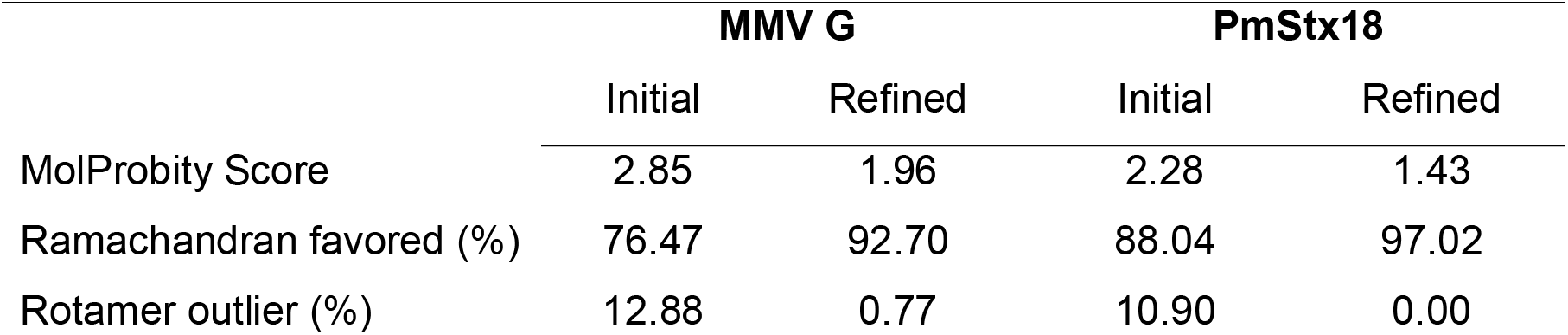
Structural refinement of the initial model of MMV G and PmStx18 predicted by the I-TASSER web server.

|  | MMV G |  | PmStx18 |  |
| --- | --- | --- | --- | --- |
|  | Initial | Refined | Initial | Refined |
| MolProbity Score | 2.85 | 1.96 | 2.28 | 1.43 |
| Ramachandran favored (%) | 76.47 | 92.70 | 88.04 | 97.02 |
| Rotamer outlier (%) | 12.88 | 0.77 | 10.90 | 0.00 |

**Figure 2.**
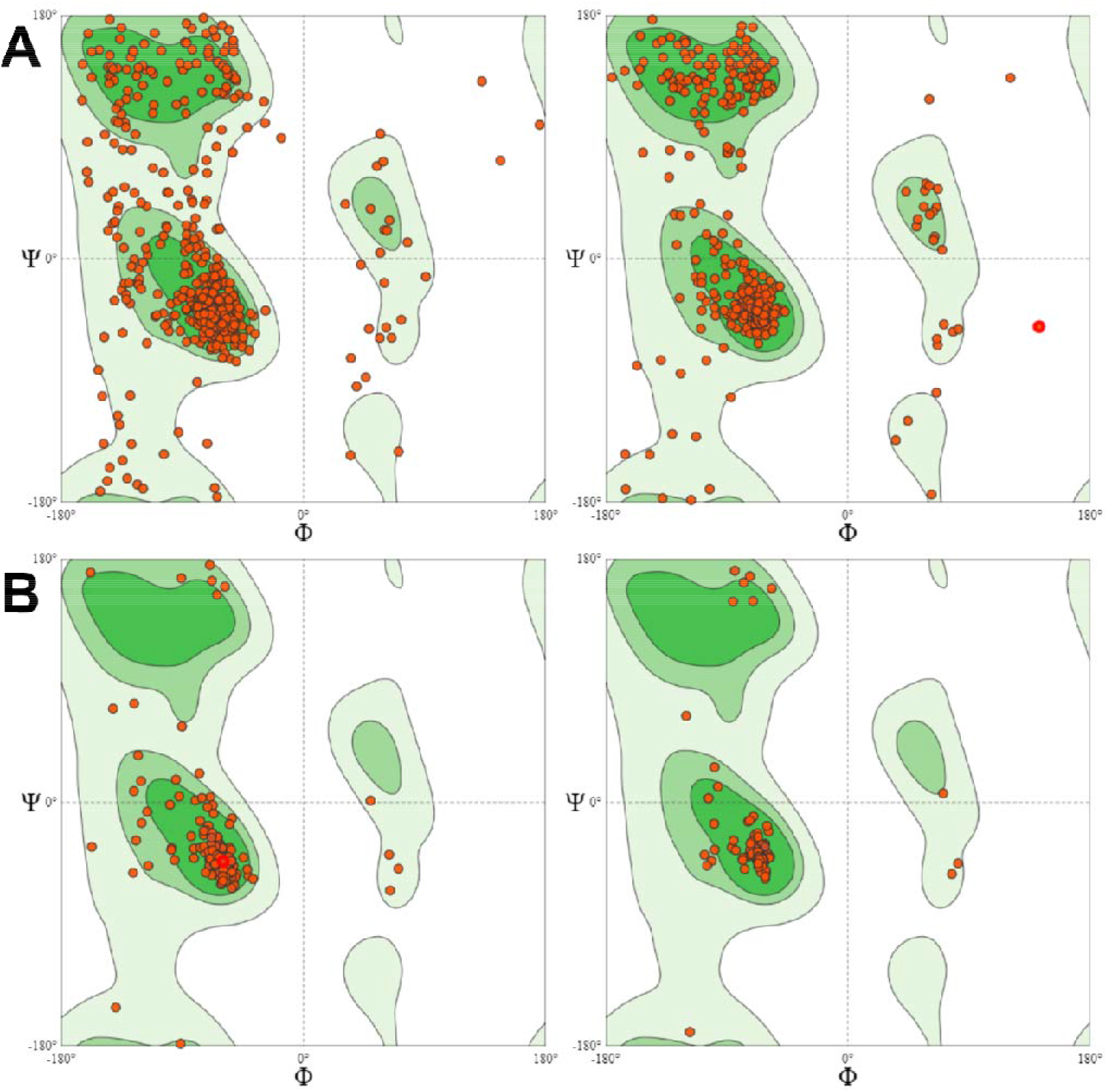
Ramachandran plots of the (A) MMV G and (B) PmSTX models (left) before and (right) after structural refinement.

### Protein-protein Docking

Various protein-protein docking web servers were used to predict the possible binding pose of MMV G with PmStx18 (Table 2). Although HADDOCK predicted a complex with a higher absolute Δ^i^G, ZDOCK complex was chosen for further experiments due to its relatively high absolute Δ^i^G and the number of interactions at its interface. ZDOCK complex ranked 2^nd^ in the highest Δ^i^G, 2^nd^ in the highest number of hydrogen bonding, 2^nd^ in the highest number of salt bridges, and 1^st^ in the highest number of vdW interaction. The 3D conformation of ZDOCK complex is shown in Figure 3, together with the respective residues present at the interface (MMV G – violet; PmStx18 – orange). The specific interaction of MMV G residues with PmStx18 residues are listed in Table 3 with the format MMV G: PmStx18. Notable MMV G residues with multiple interactions are C71, R87, W175, K233, and D279.

**Table 2.** Characterization of the predicted MMVG-PmStx18 complex by different protein-protein docking web servers.

| Docking Server | $\Delta^iG$, kcal/mol | Hydrogen bonds | Salt Bridges | Pi-stacking | vdW interaction |
| --- | --- | --- | --- | --- | --- |
| HADDOCK | -33.4 | 2 | 1 | 4 | 21 |
| ZDOCK | -31.0 | 9 | 2 | 1 | 35 |
| VakserLab | -30.8 | 8 | 1 | 0 | 34 |
| PatchDock | -29.6 | 15 | 0 | 0 | 21 |
| LZerd | -29.1 | 6 | 0 | 1 | 22 |
| InterEvDock | -22.9 | 0 | 1 | 0 | 17 |
| ClusPro | -20.3 | 5 | 3 | 1 | 14 |
| pyDockWEB | -18.8 | 0 | 0 | 1 | 25 |
| HDOCK | -16.9 | 5 | 2 | 0 | 13 |

**Figure 3.**
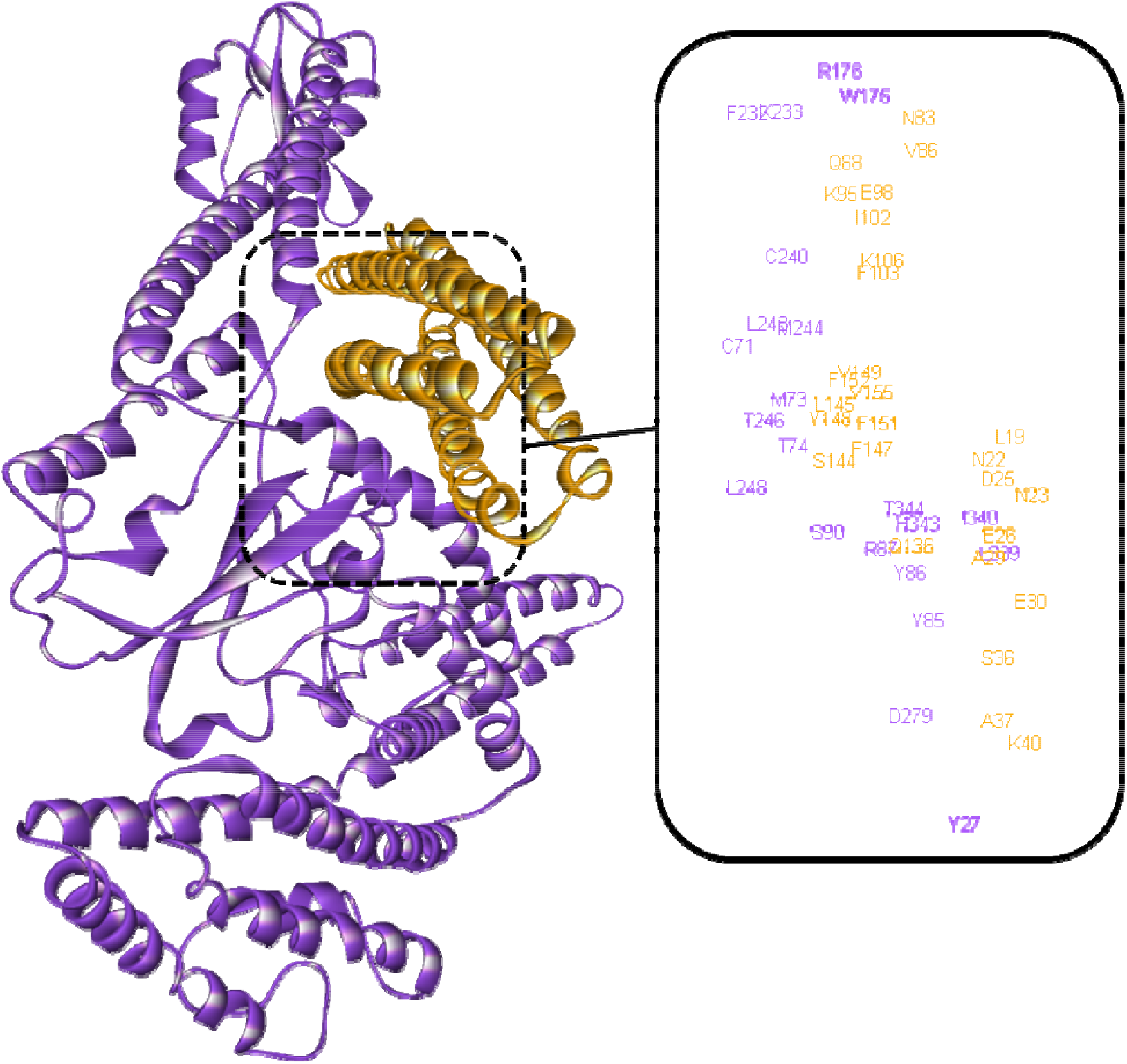
The 3D structure of MMV G-PmStx18 as predicted by the ZDOCK webserver showing the residues interacting at the interface (MMV G – violet; PmStx18 – orange).

**Table 3.** Interacting residues at the interface of the predicted MMVG-PmStx18 complex using ZDOCK web server (Format = MMVG: PmStx18). The underlined MMV G residues means that it interacts with more than one type of interaction.

| Hydrogen Bonds | Pi-Stack | Salt Bridges | vdW interaction |
| --- | --- | --- | --- |
| <u>C71</u>: F152 | H343: F147 | <u>R87</u>: D25 | <u>C71</u>: Y148 |
| Y85: E30 |  | <u>K233</u>: E98 | M73: F151, F152, V155 |
| <u>R87</u>: N22, D25, E26 |  |  | T74: F151 |
| <u>W175</u>: V86 |  |  | Y86: A29 |
| <u>K233</u>: E98 |  |  | <u>R87</u>: N22 |
| <u>D279</u>: K40, Q136 |  |  | M90: S144 |
|  |  |  | <u>W175</u>: N83 |
|  |  |  | R176: Q88 |
|  |  |  | F232: K95 (2x) |
|  |  |  | <u>K233</u>: E98 |
|  |  |  | C240: I102 (3x), Y148 |
|  |  |  | L243: T148, F103 (2x) |
|  |  |  | M244: K106, L145, V149 |
|  |  |  | T246: S144 |
|  |  |  | L248: F147 |
|  |  |  | <u>D279</u>: S36, A37 |
|  |  |  | L339: N23, E26 (2x) |
|  |  |  | I340: L19, N23 |
|  |  |  | T344: F151 (2x) |

### Molecular dynamics simulation

A 25-ns molecular dynamics simulation was conducted to investigate the changes in the conformations of the complex. Figure 4 shows how the MMV G and PmStx18 proteins changed in structure after the simulation. The complex remained intact throughout the simulation with minor conformational and structural changes in the site of attachment. Major changes in the structure of MMV G happened in the outer regions where contact with the solvent molecules is greater. The initial hydrogen bond interaction (Table 3) distances were monitored and plotted versus time (Figure 5). The analysis revealed that the hydrogen bond between R87 of MMV G and D25 of PmSTX18 remained intact after 25 ns. This suggests that R87 plays an important role in the binding affinity as additional to the high binding affinity contributed by hydrophobic interactions. Although not as intact as R87:D25, the hydrogen bond D279:K40 and W175:V86 are also notable as key interactions at the early stages of the simulation.

**Figure 4.**
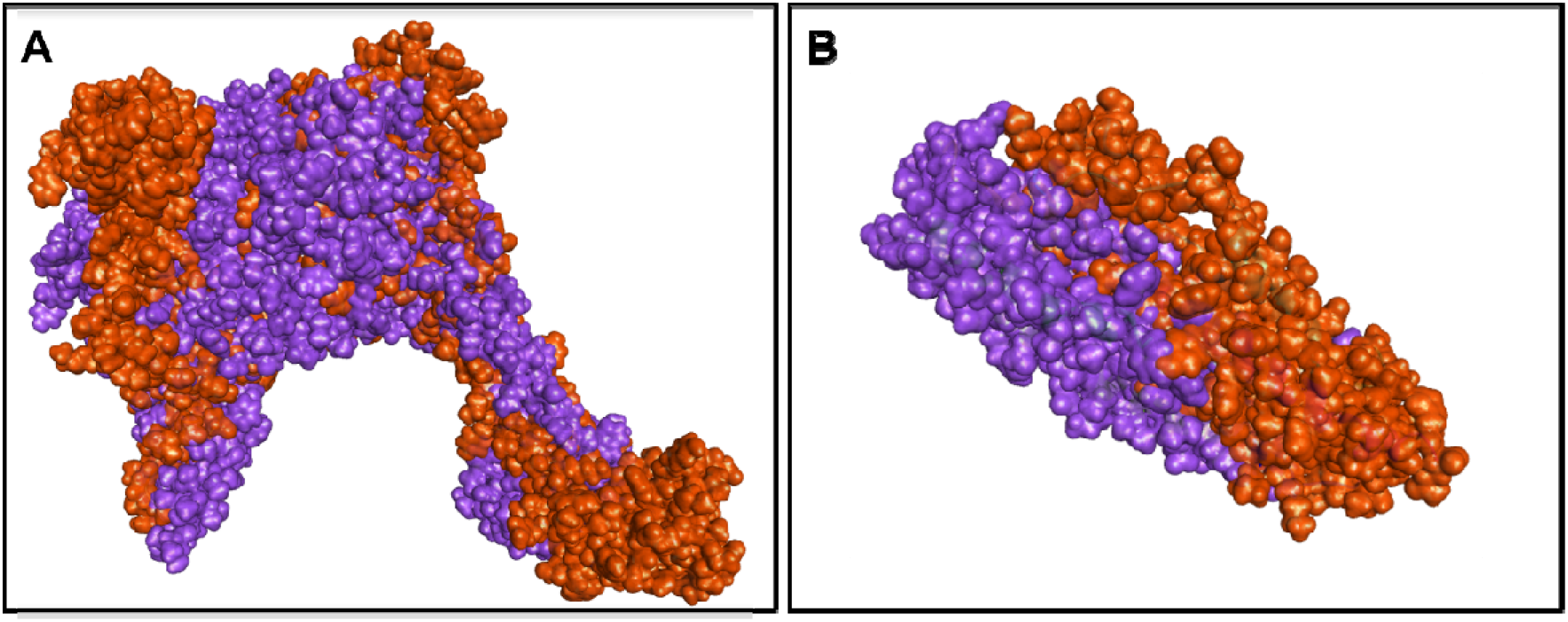
Superimposition of (A) MMV G and (B) PmStx18 chains before (violet) and after (orange) the molecular dynamics simulation.

**Figure 5.**
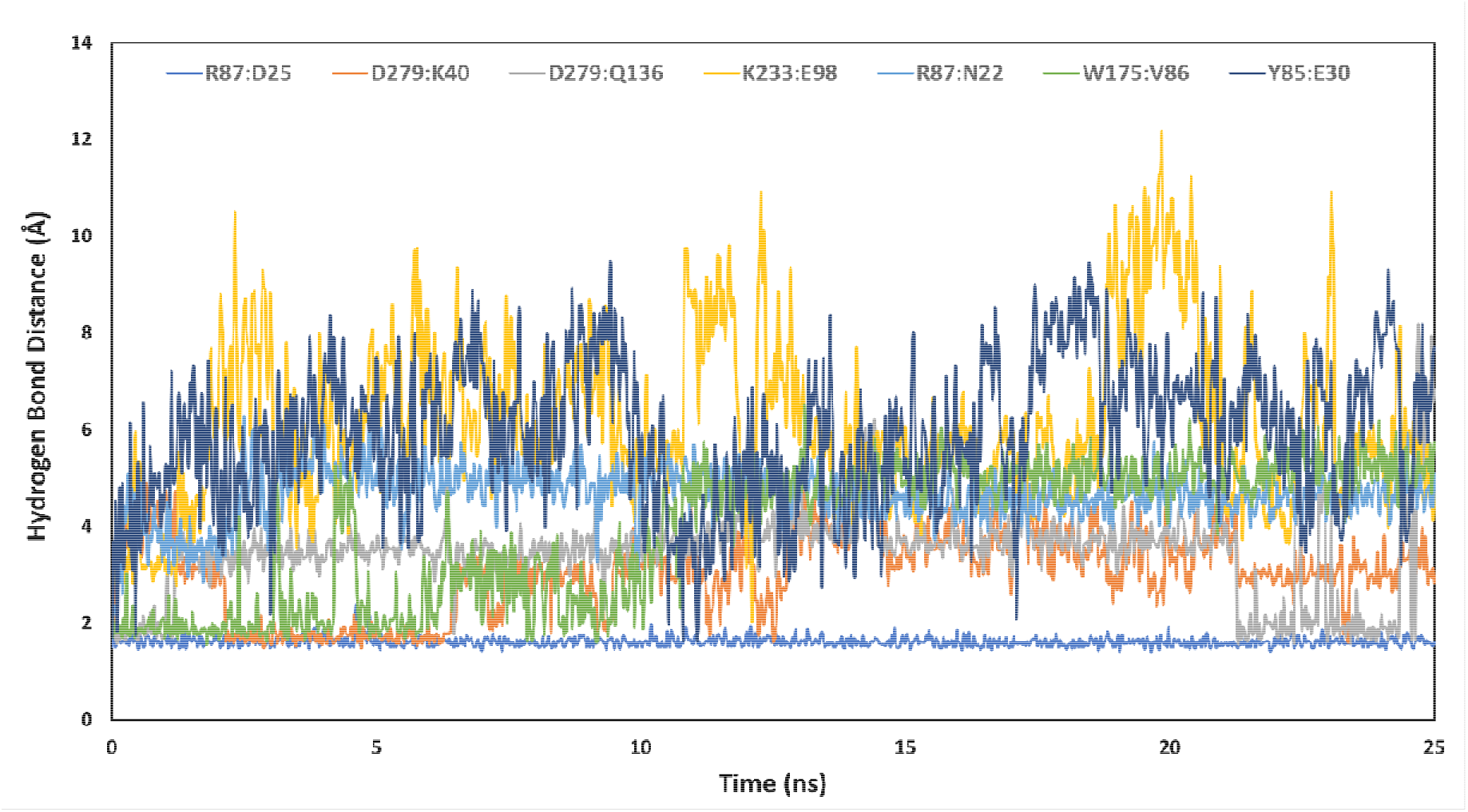
Monitoring of the initial hydrogen bond distances throughout the course of the simulation.

The interaction of MMV G with the putative PmSTX18 receptor could be a possible mechanism of entry of viral particles into host cells and subsequent infection and dissemination within the insect. This is further supported by previous studies which reported that STX18 of host cells is involved in mediating infection, such that of the *Bovine papillomavirus type 1* (BVP1) where the said receptor interacts with the viral capsid protein to facilitate infection (Bossis et al., 2005; Laniosz et al., 2007).

Viral glycoprotein interaction with cellular receptors has been a common mechanism of entry and infection for rhabdoviruses such as *Rabies virus* (RABV) and *Vesicular stomatitis virus* (VSV), prototype of genera *Lyssavirus* and *Vesiculovirus*, respectively (Belot et al., 2019). Additionally, RABV, VSV, the *Australian bat lyssavirus* (ABLV) as well as the rhabdoviral fish pathogen, *Infectious hematopoietic necrosis virus* (IHNV) infect host cells through the clathrin-mediated endocytosis (CME) pathway (Guo et al., 2019; Liu et al., 2011; Sun et al., 2005; Weir et al., 2014). In connection, this study suggests a possible mechanism for the entry and infection of MMV through the interaction of the viral glycoprotein with the putative receptor PmSTX18 since it could be a similarity shared across studied species of Rhabdoviridae. Moreover, this is the first in silico analysis performed for MMV G-PmSTX18 interaction thus, the findings of this study will contribute significantly to future studies regarding PmSTX18.

## CONCLUSION

In this study, computational and molecular interaction docking tools were used to compare the strength of the predicted protein-protein interaction of PmStx18 and MMV G. Also, molecular dynamics and simulations of the best docked protein-ligand structures revealed the dynamics information of their stability in the biological system. Based on our findings, we believe that the interaction model between the viral glycoprotein and insect vector SNARE protein can be a valuable initial step for developing a novel target specific bioinsecticide against the insect pest. Disrupting the structure stability may lead to inhibition of viral movement inside the host, which in response would restrict viral transmission to a healthy plant host.

## REFERENCES

Alviar, K. B., Rotenberg, D., Martin, K. M., & Whitfield, A. E. (2022). Identification of interacting proteins of maize mosaic virus glycoprotein in its vector, *Peregrinus maidis*. BioRxiv, 2022.02.01.478665. https://doi.org/10.1101/2022.02.01.478665

Artimo, P., Jonnalagedda, M., Arnold, K., Baratin, D., Csardi, G., De Castro, E., Duvaud, S., Flegel, V., Fortier, A., & Gasteiger, E. (2012). ExPASy: SIB bioinformatics resource portal. Nucleic Acids Research, 40(W1), W597–W603.

Barandoc-Alviar, K., Ramirez, G. M., Rotenberg, D., & Whitfield, A. E. (2017). Analysis of Acquisition and Titer of Maize Mosaic Rhabdovirus in Its Vector, Peregrinus maidis (Hemiptera□: Delphacidae). 16(September), 10–17. https://doi.org/10.1093/jisesa/iev154

Belot, L., Albertini, A., & Gaudin, Y. (2019). Structural and cellular biology of rhabdovirus entry.

Bossis, I., Roden, R. B. S., Gambhira, R., Yang, R., Tagaya, M., Howley, P. M., & Meneses, P. I. (2005a). Interaction of tSNARE syntaxin 18 with the papillomavirus minor capsid protein mediates infection. Journal of Virology, 79(11), 6723–6731.

Bossis, I., Roden, R. B. S., Gambhira, R., Yang, R., Tagaya, M., Howley, P. M., & Meneses, P. I. (2005b). Interaction of tSNARE Syntaxin 18 with the Papillomavirus Minor Capsid Protein Mediates Infection. Journal of Virology, 79(11), 6723–6731. https://doi.org/10.1128/jvi.79.11.6723-6731.2005

Bowers, K. J., Chow, D. E., Xu, H., Dror, R. O., Eastwood, M. P., Gregersen, B. A., Klepeis, J. L., Kolossvary, I., Moraes, M. A., & Sacerdoti, F. D. (2006). Scalable algorithms for molecular dynamics simulations on commodity clusters. SC’06: Proceedings of the 2006 ACM/IEEE Conference on Supercomputing, 43.

Chen, V. B., Arendall, W. B., Headd, J. J., Keedy, D. A., Immormino, R. M., Kapral, G. J., Murray, L. W., Richardson, J. S., & Richardson, D. C. (2010). MolProbity: all-atom structure validation for macromolecular crystallography. Acta Crystallographica Section D: Biological Crystallography, 66(1), 12–21.

Christoffer, C., Chen, S., Bharadwaj, V., Aderinwale, T., Kumar, V., Hormati, M., & Kihara, D. (2021). LZerD webserver for pairwise and multiple protein–protein docking. Nucleic Acids Research.

De Vries, S. J., Van Dijk, M., & Bonvin, A. M. J. J. (2010). The HADDOCK web server for data-driven biomolecular docking. Nature Protocols, 5(5), 883–897.

Dietzgen, R. G., Mann, K. S., & Johnson, K. N. (2016). Plant Virus – Insect Vector Interactions□: Current and Potential Future Research Directions. 1–21. https://doi.org/10.3390/v8110303

Fasshauer, D., Sutton, R. B., Brunger, A. T., & Jahn, R. (1998). Conserved structural features of the synaptic fusion complex: SNARE proteins reclassified as Q-and R-SNAREs. Proceedings of the National Academy of Sciences, 95(26), 15781–15786.

Guo, Y., Duan, M., Wang, X., Gao, J., Guan, Z., & Zhang, M. (2019). Early events in rabies virus infection—Attachment, entry, and intracellular trafficking. In Virus Research (Vol. 263, pp. 217–225). Elsevier B.V. https://doi.org/10.1016/j.virusres.2019.02.006

Hatsuzawa, K., Hirose, H., Tani, K., Yamamoto, A., Scheller, R. H., & Tagaya, M. (2000). Syntaxin 18, a SNAP receptor that functions in the endoplasmic reticulum, intermediate compartment, and cis-Golgi vesicle trafficking. Journal of Biological Chemistry, 275(18), 13713–13720.

Heo, L., Lee, H., & Seok, C. (2016). GalaxyRefineComplex: Refinement of protein-protein complex model structures driven by interface repacking. Scientific Reports, 6(1), 1–10.

Jiménez-García, B., Pons, C., & Fernández-Recio, J. (2013). pyDockWEB: a web server for rigid-body protein–protein docking using electrostatics and desolvation scoring. Bioinformatics, 29(13), 1698–1699.

Jourdan-Ruf, C., Marchand, J.-L., Pham, H., Markham, P., & Buduca, C. (1995). Maize streak, maize stripe and maize mosaic virus diseases in the tropics (Africa and islands in the Indian Ocean). Agriculture et Développement (Montpellier), DEC, 55–69.

Kannan, M., Ismail, I., & Bunawan, H. (2018). Maize dwarf mosaic virus: From genome to disease management. Viruses, 10(9), 492.

Karavina, C. (2014). Maize streak virus: A review of pathogen occurrence, biology and management options for smallholder farmers. African Journal of Agricultural Research, 9(36), 2736–2742.

Klobasa, W., Chu, F.-C., Huot, O., Grubbs, N., Rotenberg, D., Whitfield, A. E., & Lorenzen, M. D. (2021). Microinjection of Corn Planthopper, Peregrinus maidis, Embryos for CRISPR/Cas9 Genome Editing. Journal of Visualized Experiments: Jove, 169.

Ko, J., Park, H., Heo, L., & Seok, C. (2012). GalaxyWEB server for protein structure prediction and refinement. Nucleic Acids Research, 40(W1), W294–W297.

Kozakov, D., Hall, D. R., Xia, B., Porter, K. A., Padhorny, D., Yueh, C., Beglov, D., & Vajda, S. (2017). The ClusPro web server for protein–protein docking. Nature Protocols, 12(2), 255–278.

Krissinel, E., & Henrick, K. (2007). Inference of macromolecular assemblies from crystalline state. Journal of Molecular Biology, 372(3), 774–797.

Laniosz, V., Nguyen, K. C., & Meneses, P. I. (2007). Bovine Papillomavirus Type 1 Infection Is Mediated by SNARE Syntaxin 18. Journal of Virology, 81(14), 7435–7448. https://doi.org/10.1128/jvi.00571-07

Liu, H., Liu, Y., Liu, S., Pang, D.-W., & Xiao, G. (2011). Clathrin-Mediated Endocytosis in Living Host Cells Visualized through Quantum Dot Labeling of Infectious Hematopoietic Necrosis Virus. Journal of Virology, 85(13), 6252–6262. https://doi.org/10.1128/jvi.00109-11

Lu, H., Zhou, Q., He, J., Jiang, Z., Peng, C., Tong, R., & Shi, J. (2020). Recent advances in the development of protein–protein interactions modulators: mechanisms and clinical trials. Signal Transduction and Targeted Therapy, 5(1), 1–23.

Mueller, D. S., Wise, K. A., Sisson, A. J., Allen, T. W., Bergstrom, G. C., Bosley, D. B., Bradley, C. A., Broders, K. D., Byamukama, E., & Chilvers, M. I. (2016). Corn yield loss estimates due to diseases in the United States and Ontario, Canada from 2012 to 2015. Plant Health Progress, 17(3), 211–222.

Pierce, B. G., Wiehe, K., Hwang, H., Kim, B.-H., Vreven, T., & Weng, Z. (2014). ZDOCK server: interactive docking prediction of protein–protein complexes and symmetric multimers. Bioinformatics, 30(12), 1771–1773.

Quignot, C., Rey, J., Yu, J., Tufféry, P., Guerois, R., & Andreani, J. (2018). InterEvDock2: an expanded server for protein docking using evolutionary and biological information from homology models and multimeric inputs. Nucleic Acids Research, 46(W1), W408–W416.

Rao, V. S., Srinivas, K., Sujini, G. N., & Kumar, G. N. (2014). Protein-protein interaction detection: methods and analysis. International Journal of Proteomics, 2014.

Roy, A., Kucukural, A., & Zhang, Y. (2010). I-TASSER: a unified platform for automated protein structure and function prediction. Nature Protocols, 5(4), 725–738.

Schneidman-Duhovny, D., Inbar, Y., Nussinov, R., & Wolfson, H. J. (2005). Geometry-based flexible and symmetric protein docking. Proteins: Structure, Function, and Bioinformatics, 60(2), 224–231.

Sun, X., Yau, V. K., Briggs, B. J., & Whittaker, G. R. (2005). Role of clathrin-mediated endocytosis during vesicular stomatitis virus entry into host cells. Virology, 338(1), 53–60. https://doi.org/10.1016/j.virol.2005.05.006

Tovchigrechko, A., & Vakser, I. A. (2006). GRAMM-X public web server for protein–protein docking. Nucleic Acids Research, 34(suppl_2), W310–W314.

Weir, D. L., Laing, E. D., Smith, I. L., Wang, L. F., & Broder, C. C. (2014). Host cell virus entry mediated by Australian bat lyssavirus G envelope glycoprotein occurs through a clathrin-mediated endocytic pathway that requires actin and Rab5. Virology Journal, 11(1). https://doi.org/10.1186/1743-422X-11-40

Yan, Y., Tao, H., He, J., & Huang, S.-Y. (2020). The HDOCK server for integrated protein–protein docking. Nature Protocols, 15(5), 1829–1852.

Yang, J., Yan, R., Roy, A., Xu, D., Poisson, J., & Zhang, Y. (2015). The I-TASSER Suite: protein structure and function prediction. Nature Methods, 12(1), 7–8.

Yao, J., Rotenberg, D., Afsharifar, A., Barandoc-Alviar, K., & Whitfield, A. E. (2013). Development of RNAi methods for Peregrinus maidis, the corn planthopper. PloS One, 8(8), e70243.

Yoon, T.-Y., & Munson, M. (2018). SNARE complex assembly and disassembly. Current Biology, 28(8), R397–R401.

Zhang, Y. (2008). I-TASSER server for protein 3D structure prediction. BMC Bioinformatics, 9(1), 1–8.

Zhang, Y., & Skolnick, J. (2005). TM-align: a protein structure alignment algorithm based on the TM-score. Nucleic Acids Research, 33(7), 2302–2309.

