## Supplemental Figure for "In silico structure-based analysis of the predicted protein-protein interaction of Syntaxin-18, a putative receptor of *Peregrinus maidis* Ashmead (Hemiptera: *Delphacidae*) with Maize mosaic virus glycoprotein"

**SUPPLEMENTAL FIGURES**

Appendix 1. I-TASSER prediction of the secondary structure of PmSTX18.

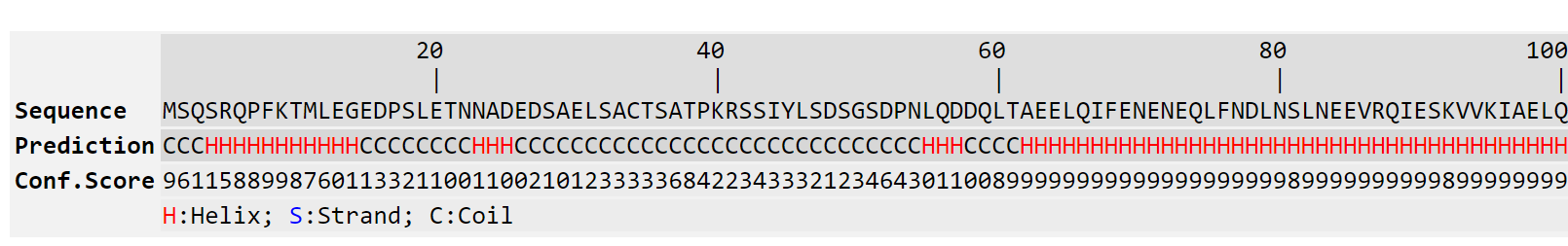

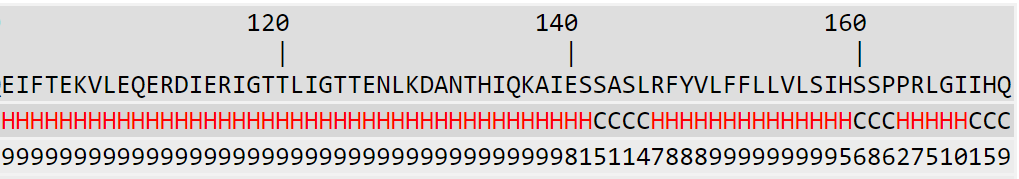

Appendix 2. I-TASSER prediction of the secondary structure of MMV G.

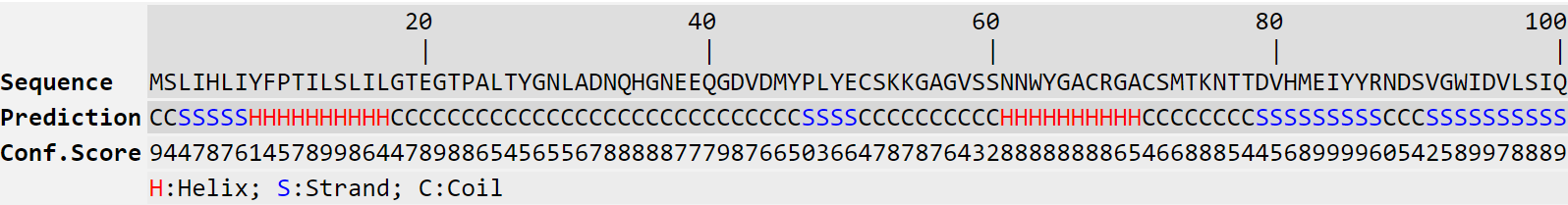

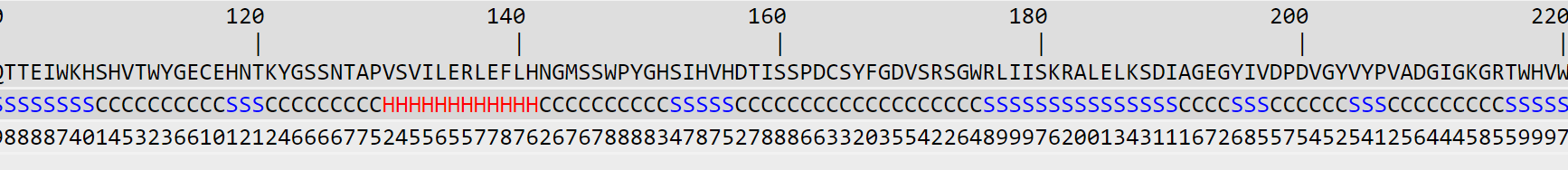

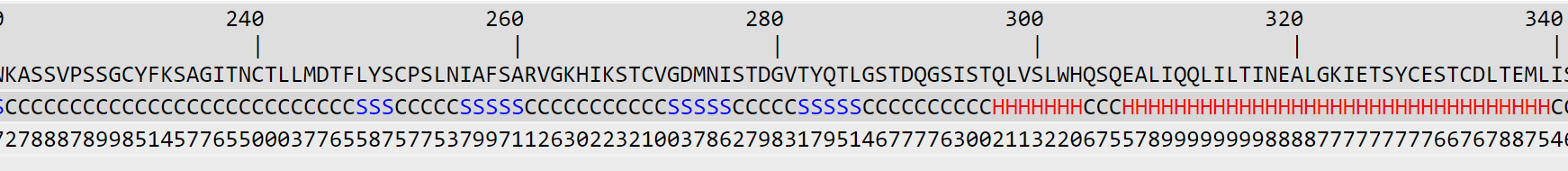

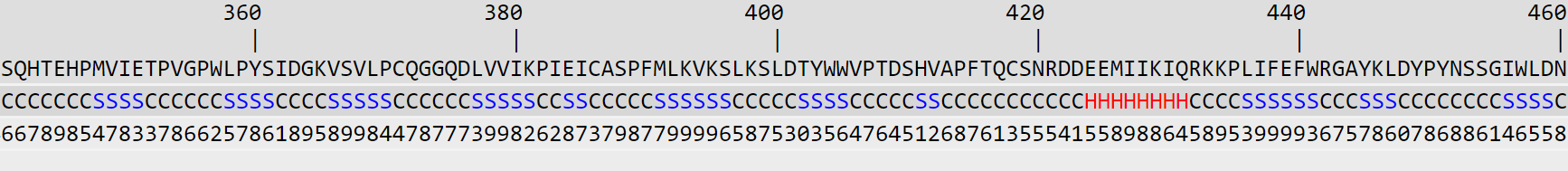

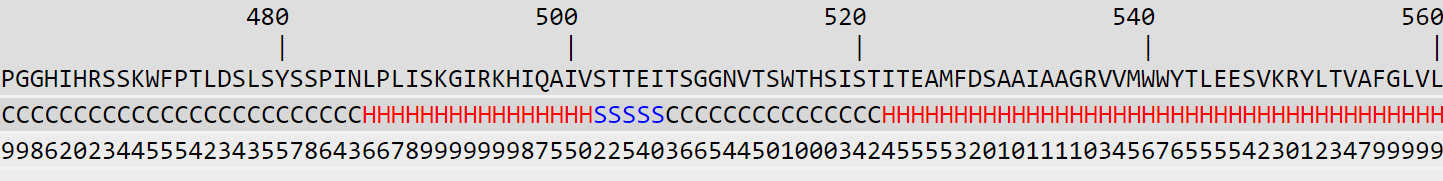

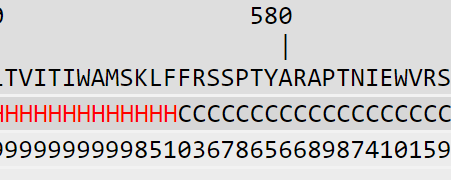
